# Hippocampal-cortical functional connectivity during memory encoding and retrieval

**DOI:** 10.1101/2022.09.02.506311

**Authors:** Liisa Raud, Markus H. Sneve, Didac Vidal-Piñeiro, Øystein Sørensen, Line Folvik, Hedda T. Ness, Athanasia M. Mowinckel, Håkon Grydeland, Kristine B. Walhovd, Anders M. Fjell

## Abstract

Memory encoding and retrieval are critical sub-processes of episodic memory. While the hippocampus is involved in both, its connectivity with the neocortex during memory processing in humans has been elusive. This is partially due to variations in demands in common memory tasks, which inevitably recruit cognitive processes other than episodic memory. Conjunctive analysis of data from different tasks with the same core elements of encoding and retrieval can reduce the intrusion of patterns related to subsidiary perceptual and cognitive processing. Leveraging data from two large-scale functional resonance imaging studies with different episodic memory tasks (514 and 237 participants), we identified core hippocampal-cortical networks active during memory processing. Anterior and posterior hippocampus had distinct connectivity profiles, which were stable across resting state and memory tasks. Whereas no encoding-specific connectome emerged across tasks, during retrieval hippocampal connectivity was increased with areas known to be active during recollection, including medial prefrontal, inferior parietal, and parahippocampal cortices. This indicates that the stable functional connectivity of the hippocampus along its longitudinal axis is superposed by increased functional connectivity with the recollection network during retrieval, while encoding connectivity likely reflects contextual factors.

## Introduction

Encoding and retrieval are critical processes of episodic memory, both of which rely on the interactions between the hippocampus (HC) and the neocortex (Eichenbaum, 2017, 2004). Encoding refers to creating a neural imprint of an original event, whereas retrieval refers to accessing that imprint at a later point in time (Tulving, 1984). Functional magnetic resonance imaging (fMRI) studies indicate activation of broad cortical networks beyond the HC during both encoding and retrieval (McCormick et al., 2010; Rugg and Vilberg, 2013; Sneve et al., 2015; Spaniol et al., 2009). However, it remains elusive to which degree these joint activations reflect memory, as opposed to supporting cognitive processes common to any cognitive task. Here, we identified the hippocampal networks of encoding and retrieval using fMRI functional connectivity in two different studies with large sample sizes. As the studies employed different episodic memory tasks that nonetheless targeted the same fundamental processes of encoding and retrieval, converging connectivity across the tasks can expose HC-cortical connectomes of episodic memory.

HC organization in humans follows a gradient along its longitudinal axis (Poppenk et al., 2013; Strange et al., 2014), as evidenced by different anatomical connectivity of the anterior and posterior hippocampus (Dalton et al., 2021; Vos de Wael et al., 2018) (aHC and pHC, respectively), gene expression (Vogel et al., 2020), volumetric changes in neuropsychiatric disorders (Geuze et al., 2005), life-span changes in micro- and macrostructure (Langnes et al., 2020; Malykhin et al., 2017), cognitive function (Grady, 2020; Kühn and Gallinat, 2014; Nadel et al., 2013; Poppenk et al., 2013; Reagh and Ranganath, 2018; Ritchey et al., 2015; Zeidman et al., 2015; Zeidman and Maguire, 2016), fMRI activity (Langnes et al., 2019; Spaniol et al., 2009), and functional connectivity during rest (Adnan et al., 2016; Blessing et al., 2016; Chase et al., 2015; Libby et al., 2012; Przeździk et al., 2019; Qin et al., 2016; Robinson et al., 2016; Vos de Wael et al., 2018; Ward et al., 2014) and encoding (Beason-Held et al., 2021; Tang et al., 2020). Whereas aHC is more active during encoding, pHC is relatively more active during retrieval (Langnes et al., 2019; Spaniol et al., 2009). Meta-analytic synthesis of terms related to the aHC and pHC also indicate a preferred association between aHC and encoding, while both aHC and pHC have been associated with retrieval (Grady, 2020). In addition, aHC is simultaneously active with the dorsal-attentional network during encoding, possibly since encoding involves externally triggered stimulus manipulation (Kim, 2015). In contrast, pHC activity coincides with the default mode network activity during retrieval, putatively reflecting introspective cognition (Kim, 2015). Although simultaneous activity does not necessarily mean increased communication, recent evidence does indicate increased synchronization, measured by fMRI functional connectivity, between the pHC and the default mode network during retrieval (Fritch et al., 2021), despite resting state connectivity studies predominantly reporting stronger connectivity between the default mode network and the aHC (Zheng et al., 2021). This framework sets predictions for distinct connectivity for aHC with externally oriented attentional networks during encoding, and pHC with the internally oriented default mode network during retrieval.

We estimated fMRI functional connectivity of the HC with the whole neocortex during encoding and retrieval in two large-scale studies (Study 1 n = 514 participants, age range 6-81 years, mean = 40, SD = 18; Study 2 n = 237 participants, age range 10-80 years, mean = 25, SD = 21) with two different episodic memory tasks (Figure 1). Both tasks included in-scanner encoding and retrieval sessions, but differed in various aspects, including task instructions, stimulus categories, and test interval (Table 1). We first tested whether the known distinction between the aHC and pHC connectivity during rest (Adnan et al., 2016; Blessing et al., 2016; Chase et al., 2015; Libby et al., 2012; Przeździk et al., 2019; Qin et al., 2016; Robinson et al., 2016; Vos de Wael et al., 2018; Ward et al., 2014) is also prevalent during memory processing. Secondly, we hypothesized specific HC-cortical connectomes for encoding and retrieval along the long-axis of the HC. Lastly, we tested the proposed relationship between the aHC and attentional networks during encoding, and pHC and the default mode network during retrieval (Fritch et al., 2021; Kim, 2015). Crucially, the conjunctive connectivity patterns across the two studies can expose HC-cortical networks relevant to encoding and retrieval, reducing the intrusion of patterns related to subsidiary perceptual and cognitive processing to the degree these differ between the tasks.

**Figure 1.**
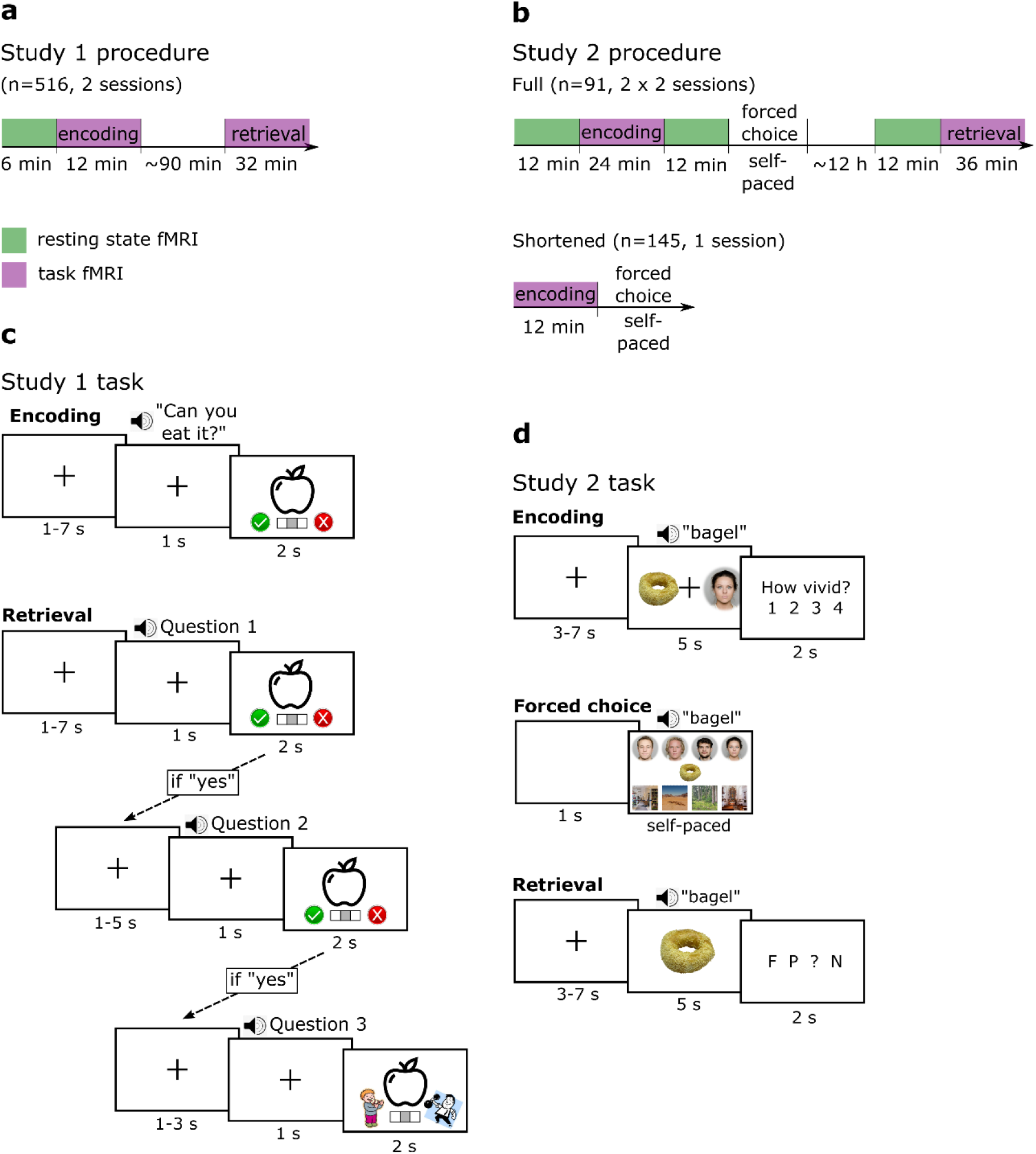
**(a-b)** Study procedures. The colored blocks represent time spent in the fMRI scanner. The encoding and retrieval tasks were performed in separate scanning sessions. In Study 2, the full procedure was repeated twice per participant, except for participants younger than 18 years old, who only completed a shortened procedure in a single session. **(c)** Memory task in Study 1. During encoding, participants had to respond “yes” or “no” to whether they can eat the item or whether they can lift it. During the surprise retrieval test, they had to answer sequentially to three questions, with the second and third presented only if the answer to the preceding question was “yes”. Question 1: “Have you seen this item before?”; question 2: “Can you remember what you were supposed to do with the item?”; Question 3: “Were you supposed to eat it or lift it?” **(d)** Memory task in Study 2. During encoding, participants were instructed to learn an association between an item and a specific face or place. Their learning was tested immediately after encoding with an eight-alternative forced choice test. During retrieval, participants were shown each item and they had to choose whether it had been associated with a face (F), with a place (P), whether they had seen the item, but couldn’t remember whether it was associated if a face or a place (?), or whether the item was new (N).

**Table 1.** Task characteristics of the two studies. *128 items per visit, with two visits for adult sample, adding up to 256 items. Participants younger than 18 years (n = 145) went through a shortened procedure in a single session.

| **Task characteristics** | **Study 1** | **Study 2** |
| --- | --- | --- |
| Encoding type | Incidental | Intentional |
| Encoding instructions | Can you eat/lift the item? | Make an association between the object and the face/place |
| Stimulus modality | Visual and auditory | Visual and auditory |
| Number of learned items | 100 | 128 (64 for shortened procedure)* |
| Intermediate retrieval | None | Forced choice after encoding |
| In-scanner retrieval type | Cued recollection | Cued recollection |
| Interval between encoding and in-scanner retrieval | 1.5 hours | 12 hours |

## Results

### Behavior

The study procedures and memory tasks are depicted in Figure 1 A-D and the major differences between the tasks are listed in Table 1. Shortly, participants in Study 1 had to make item-action evaluations, as they were asked whether they can eat (or lift) an item showed on the screen. They were tested 90 minutes later in a surprise retrieval session, testing if they remembered whether each item had been associated with ‘eating’ or ‘lifting’. Both the encoding and retrieval sessions were completed in the fMRI scanner. Participants correctly remembered the associations of 44% (sd = 19%) of the items (Figure 2), corrected for estimated guessing rates (see materials and methods for details). Participants in Study 2 were specifically instructed to form memory associations between items and faces or places during the encoding phase. Their learning performance was tested immediately after encoding with a forced choice test with eight alternatives (chance rate 12.5%), in which participants chose the correct face or place for 66% of the items (sd = 18%). A recollection test followed 12 hours later, in which participants correctly indicated associations with face or place in 59% (sd = 24%; corrected for guessing rates) of the previously learned items. Encoding and final recollection test were completed in the fMRI scanner, whereas the immediate forced choice test was done outside of the scanner. Note that participants below 18 years of age went through a shortened protocol without the final retrieval and resting state data. In adults, the performance in the forced choice and the retrieval test correlated strongly with r = 0.95.

**Figure 2.**
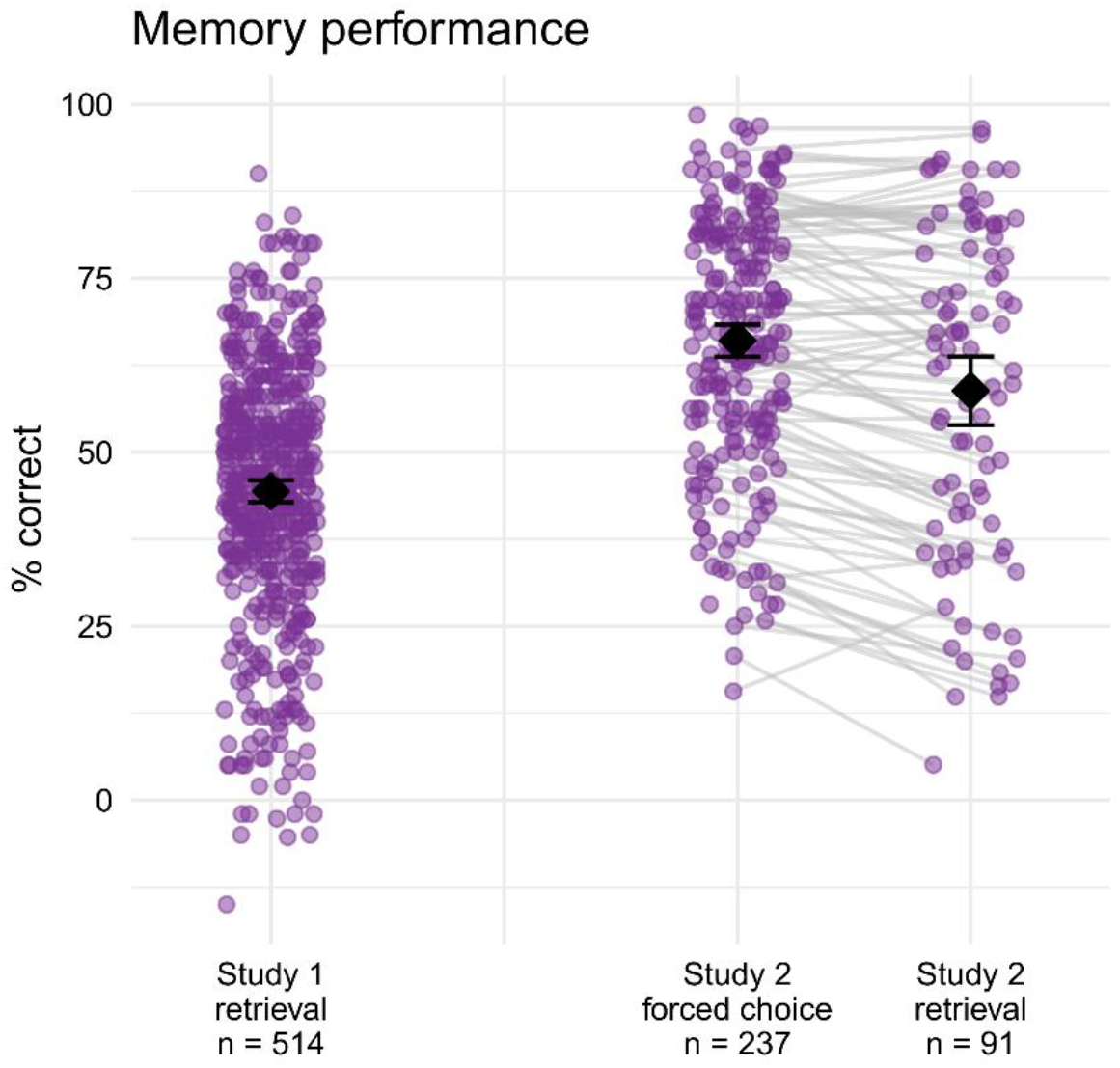
Associative memory performance as a percentage of correct answers from all items. Note that the retrieval performance is corrected for estimated guessing rates in both studies, resulting in negative values for some participants. The purple dots indicate each participant, the lines connect the participants in each test who went through the full procedure in Study 2, the black diamonds represent the sample means, and error bars represent 95% confidence intervals.

### Stable hippocampal functional connectivity during resting state and memory tasks

fMRI was recorded during encoding and retrieval phases in both studies, together with additional resting state measurement for most participants (Figure 1a-b). The HC was defined based on the automatic subcortical segmentation in FreeSurfer and further divided into aHC and pHC segments based on the MNI coordinate of y = −21, which marks the approximate position of uncus in the parahippocampal gyrus (Poppenk et al., 2013). The cortex was divided into 400 regions based on the Schaefer atlas (Schaefer et al., 2018). For the resting state, functional connectivity of the two HC seeds (aHC and pHC) with each cortical region was estimated by correlating their denoised BOLD time series. Task functional connectivity was calculated by correlational psychophysiological interaction analysis (cPPI; Fornito et al., 2012), which isolates connectivity occurring during specified task-periods, controlling for task-independent co-activation. The PPI variable was calculated simultaneously for all task events and trials, irrespective of the trial type and recollection outcome. As such, it represents event-related connectivity contrasted to implicit baseline during the task period and is similar to the beta-series correlation technique (Di et al., 2021). HC-cortical connectivity values were averaged across the left and right HC. The whole brain statistical analyses were performed using linear models separately for the resting state and memory tasks. First, a base linear model was estimated for each HC-cortical pair predicting connectivity from the HC segment (aHC vs pHC) while controlling for age and age^2^ to account for non-linear age effects, sex, and participant ID as a random variable. The task phase (encoding vs retrieval) was added as an additional predictor to the task data in later analyses where appropriate. Whereas significance was determined using a cut-off value of p < 0.05 (corrected for multiple comparisons using false discovery rate correction; Benjamini and Hochberg, 1995), effect sizes were used to visualize the results. Conjunction across the cognitive states and/or studies was deemed significant if all the constitute analyses had a p < 0.05 (Nichols et al., 2005). The list of all significant regions together with their central coordinates for all models are available as html-reports in the dedicated project at Open Science Framework (https://osf.io/834hz/).

The raw connectivity maps (i.e. the Fisher z-transformed (partial) correlations) of the HC during resting state and memory tasks showed widespread positive connectivity, which was strongest with the medial and lateral temporal cortices, temporal poles, medial parietal cortices, medial prefrontal, and orbitofrontal cortices (Supplementary Figure 1). The spatial patterns of the raw connectivity values were similar across the two studies (correlation coefficients of the spatial connectivity patterns of corresponding states between studies ranging between 0.74 to 0.78) as well as between the resting state, encoding, and retrieval data (correlation coefficients within each study ranging between 0.72 and 0.99).

The HC segment contrast (aHC vs pHC) in each base model was used to identify the differential connectivity patterns of the aHC and pHC (Figure 3a-b). In conjunction (Figure 3c), aHC was more strongly connected than the pHC with the temporal pole, parahippocampal, retrosplenial, orbitofrontal, and sensorimotor cortices. In contrast, pHC was more strongly connected than aHC with precunei, insulae, medial prefrontal, posterior cingulate, inferior parietal, and dorsolateral prefrontal cortices. The results were similar across both studies, as well as across the resting and task states (spatial correlations of the T-value maps for the aHC-pHC contrast ranging between 0.78 and 0.93).

**Figure 3.**
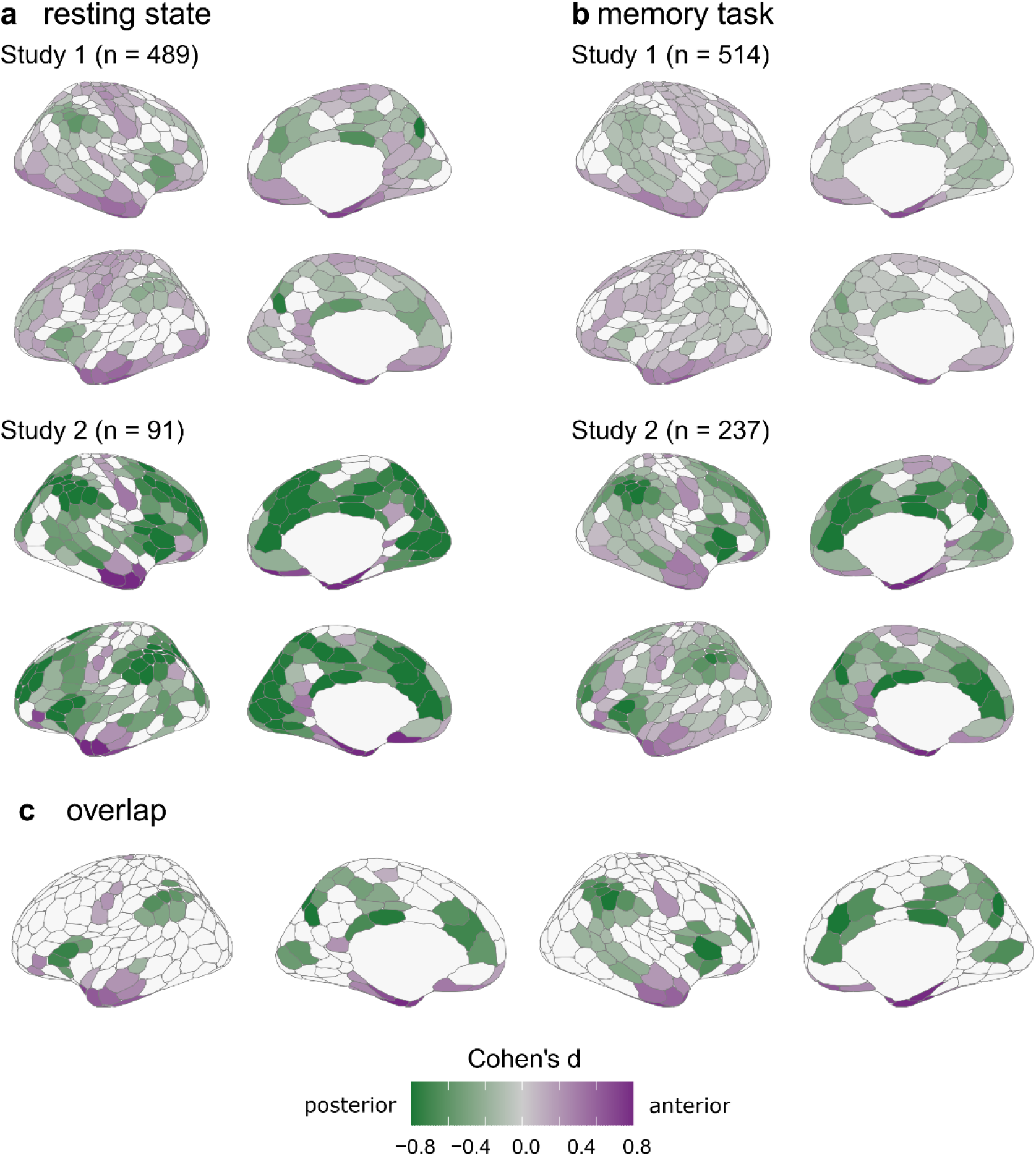
HC segment main effects during resting state (**a**) and memory tasks (**b**). The values represent average Cohen’s d effect sizes for the aHC and pHC t-contrast in linear models (controlling for sex, age, and age^2^), masked for p < 0.05 (fdr-corrected). Purple values indicate parcels where HC-cortical connectivity was stronger with the aHC than with the pHC, and green values indicate the parcels with stronger pHC than aHC connectivity. (**c**) overlap across the samples and states with the values indicating mean effect sizes across all four models in regions where all models showed a significant HC segment main effect.

In sum, the raw hippocampal connectivity patterns were similar across the resting state and memory processing, as were the distinct connectivity profiles of the aHC and pHC, in accordance with the previous studies of resting state connectivity along the longitudinal axis of the HC (Adnan et al., 2016; Blessing et al., 2016; Chase et al., 2015; Libby et al., 2012; Przeździk et al., 2019; Qin et al., 2016; Robinson et al., 2016; Vos de Wael et al., 2018; Ward et al., 2014).

### Retrieval, but not encoding, exhibited task-general hippocampal connectivity

To test whether the HC had distinct encoding- and retrieval-related connectivity profiles, the base linear models of the task data were expanded by additional task phase contrast (encoding vs retrieval), as wells as its interaction with the HC segment to test whether the task phase connectivity profiles differed between the aHC and pHC. The spatial correlation of the t-values for the encoding-retrieval contrast across the two studies was moderate with r = 0.53 (p < 0.001) and r = 0.38 (p = 0.002) for left and right hemisphere, respectively (non-parametric p-values corrected for spatial autocorrelation; Burt et al., 2020).

There was considerable overlap between the studies for retrieval-related connectivity (Figure 4). In both studies, HC had stronger connectivity during retrieval than during encoding with the anterior temporal, medial prefrontal, inferior parietal, and parahippocampal cortices. Noticeably, many of these areas are previously reported to be active during memory retrieval, sometimes referred to as the core memory recollection network (Kim, 2010; Rugg and Vilberg, 2013), and also constitute parts of the default mode network.

**Figure 4.**
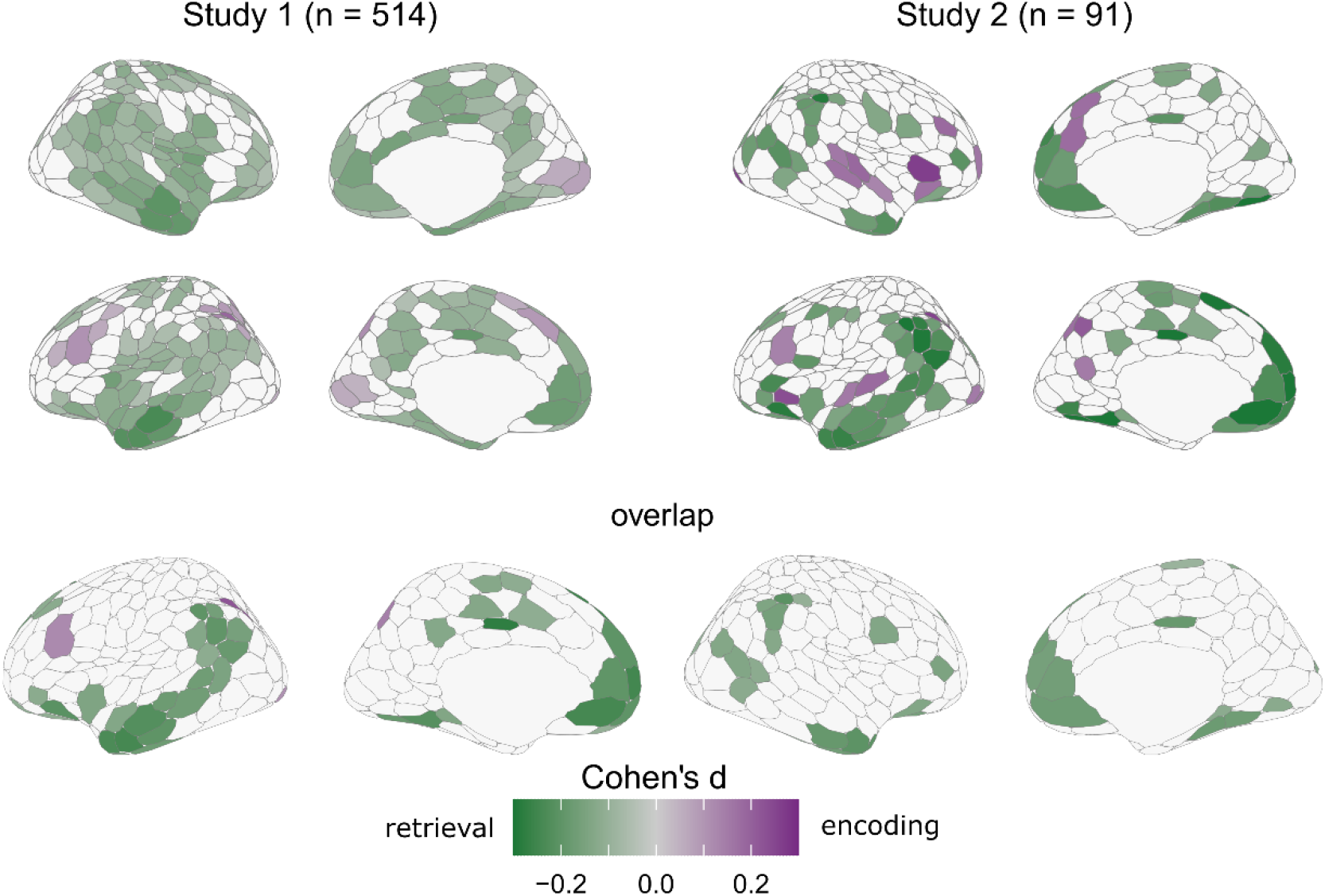
Task phase main effects in the task data. The values represent Cohen’s d’s for the encoding and retrieval t-contrast in linear models including task phase, HC segment, and their interaction as predictors (controlling for sex, age, and age^2^), masked for p < 0.05 (fdr-corrected). Purple values indicate regions in which HC-cortical connectivity was stronger during encoding than retrieval, and green values indicate the parcels with stronger retrieval than encoding connectivity. Overlap reflects average Cohen’s d, masked by significant t-values across both studies.

The overlap for increased encoding compared to retrieval connectivity across the studies included limited regions in the control network in the left hemisphere (inferior parietal sulcus and lateral prefrontal cortex). Both studies showed increased encoding connectivity also with visual cortices, though the exact regions did not overlap. In Study 1, (incidental) encoding additionally had stronger HC-cortical connectivity with superior parietal cortex, while in Study 2, encoding connectivity was stronger with insulae and auditory cortices, right medial prefrontal cortex, left precuneus, and retrosplenial cortex.

There were no significant interactions of the task-phase and HC segment with any of the cortical regions in Study 1 and only with a few isolated regions in Study 2 (see supplementary reports at Open Science Framework for detailed lists of significant regions in each study), indicating a lack of functional dissociation between the aHC and pHC connectivity with respect to encoding and retrieval.

In sum, connectivity differences between encoding and retrieval were affected by task demands. Both studies showed stronger HC connectivity with recollection-related areas. In contrast, conjunction for encoding-related connectivity across studies was limited, potentially due to different task instructions (incidental vs intentional encoding) and stimulus categories.

### Limited evidence for the encoding-aHC-attentional and retrieval-pHC-default mode network associations

Given previous meta-analytic evidence for co-activation of the aHC with the dorsal-attentional network during encoding (Kim, 2015), we tested the HC segment and cognitive state differences in HC connectivity with known attentional networks. In addition to the dorsal-attentional network, we also included other networks related to external processing and increased cognitive demands, i.e. the salience and frontoparietal control networks. Functional connectivity with HC was averaged across regions belonging to each network and differences in the HC-network connectivity were tested using mixed-effects linear models with planned contrasts: HC segment (aHC vs pHC), cognitive state (two orthogonal contrasts: resting vs task state; encoding vs retrieval), and their interactions, controlling for age, age^2^, and sex. The test statistics together with the p-values for statistical inference are listed in Table 2. However, the discussion of the results is guided by effect sizes measured by Cohen’s d (in the following presented in the format *S1d = x, S2d = y*, for Study 1 and Study 2, respectively), for which 0.2 is generally considered a small effect (Cohen, 1992). To prevent discarding very small effects with significant p-values a negligible, we will consider an effect size larger than 0.15 in at least one of the studies relevant, given that the direction of the effects agreed across the studies.

**Table 2.**
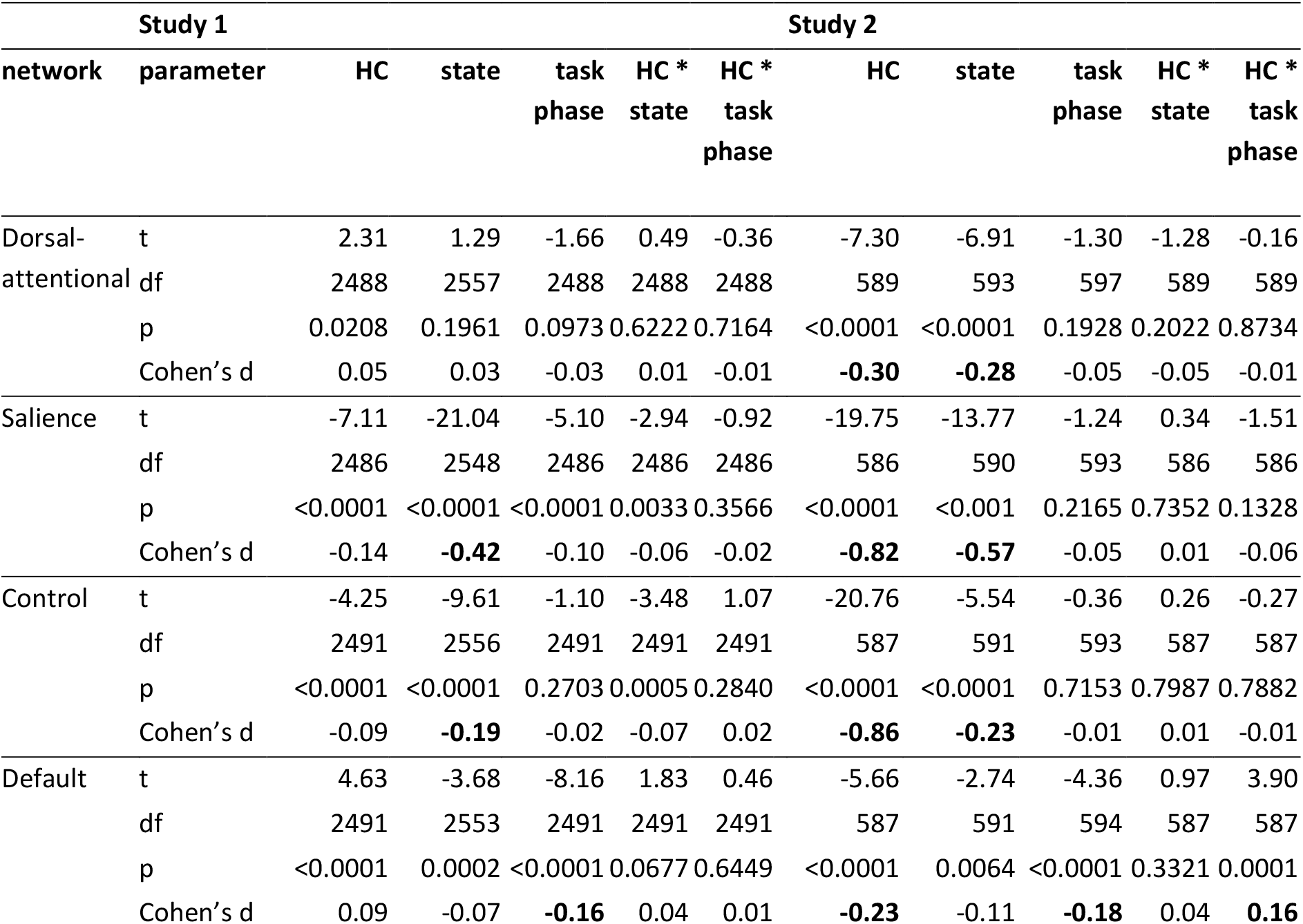
Statistical results for planned contrasts for HC connectivity with cortical networks. Effect sizes of d >= 0.15 marked in bold. The HC t-values represent the contrast aHC > pHC, the state represents rest > task, and the task phase represents encoding > retrieval.

In contrast to our predictions, the pHC had stronger connectivity with the externally oriented networks than the aHC (Figure 5b). This effect was prominent for the salience (S1d = −0.14, S2d = −0.82; negative values indicate larger pHC than aHC connectivity, and vice versa for positive values) and control networks (S1d = −0.09, S2d = −0.86), whereas results diverged across the studies for the dorsal-attentional network (S1d = 0.05, S2d = −0.30). Connectivity was stronger during the task than during the rest for the salience (S1d = −0.42, S2d = −0.57; negative values indicate stronger task than rest connectivity) and control networks (S1d = −0.19, S2d = −0.23); and for dorsal-attentional network in Study 2 (S1 d = 0.02, S2d = −0.28). Differences between encoding and retrieval states as well as all interaction effect sizes were negligible.

**Figure 5.**
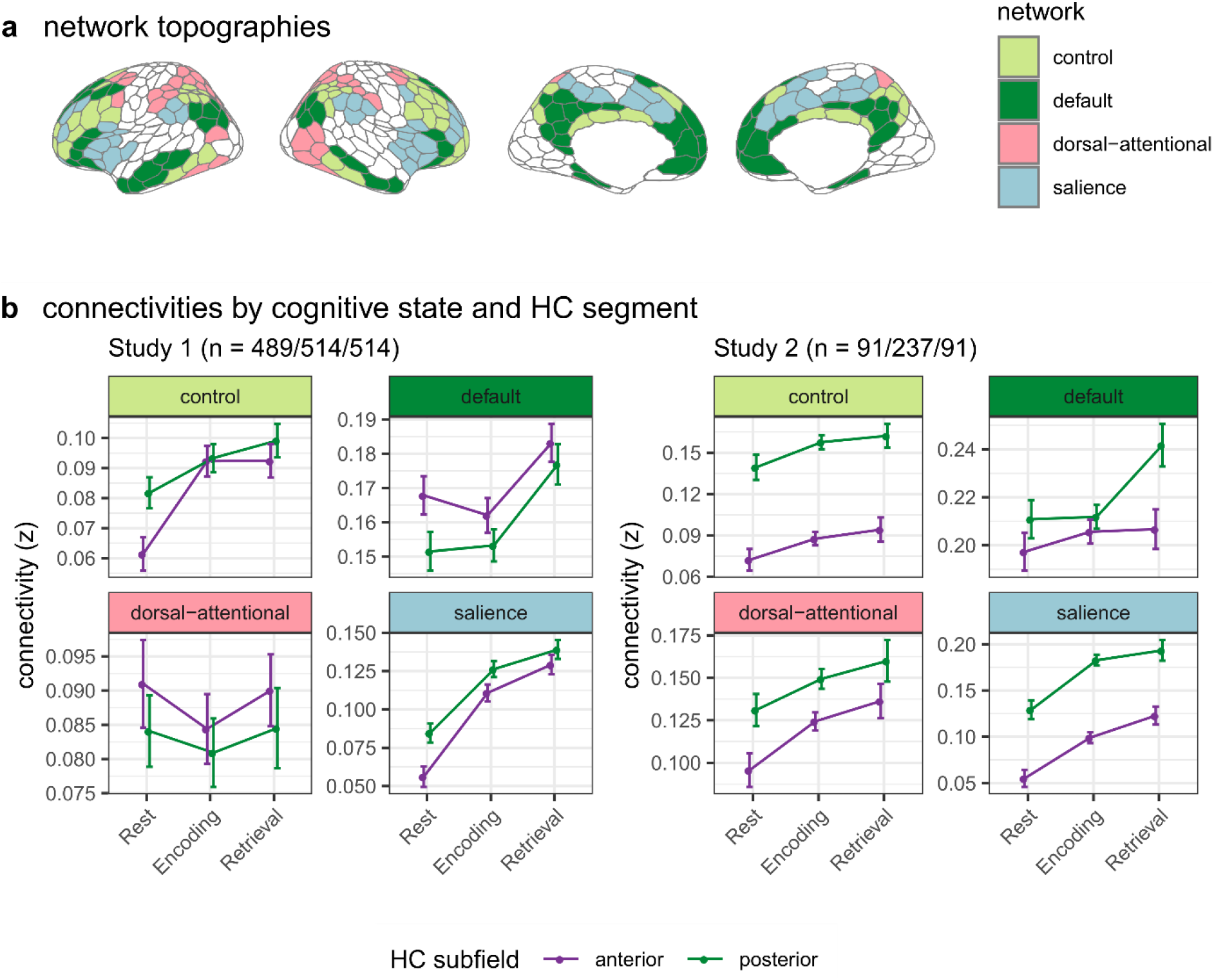
**(a)** Resting state network topologies according to Schaefer et al. (2018) **(b)** Averaged functional connectivities with the HC by cognitive state and HC segment. The error bars represent 95% confidence intervals, corrected for repeated measurements. The sample sizes in brackets refer to rest/encoding/retrieval. HC = hippocampus.

The connectivity differences between aHC and pHC with the default mode network were in different directions across the studies (S1d = 0.09, S2d = −0.23). Both studies showed stronger overall HC connectivity during retrieval than during encoding (HC main effect S1d = −0.16, S2d = −0.18; negative values indicate stronger retrieval than encoding connectivity), and Study 2 additionally showed an interaction between task phase and HC segment (S1d = 0.01, S2d = 0.16), with connectivity increase with pHC during retrieval.

In sum, our results are against encoding-specific upregulation of aHC connectivity with externally oriented networks, although increased task connectivity (compared to resting state) with the control and salience networks was observed for the whole HC. Instead, pHC had stronger connectivity than aHC with salience and frontoparietal control networks. In addition, there was support for upregulation of the internally oriented default mode network with HC during retrieval, and particularly with the pHC after a 12 hour retention period in Study 2.

### No associations between hippocampal connectivity and memory

To test the associations between HC-cortical connectivity and memory performance, the whole-brain linear models were extended with task performance as a predictor variable. No regions showed significant memory main effects in neither resting nor task data. However, due to interaction effects between memory performance and HC segment in the task data (Supplementary Figure 2), the models were run separately for the anterior and posterior HC. Initial analysis indicated that in Study 1, higher connectivity with lateral and medial temporal lobe, insula, and inferior frontal cortex was associated with lower memory performance, which was stronger and more widespread for the pHC than the aHC (Supplementary Figure 3). However, these effects disappeared after additional control analyses, taking into account that people with lower memory performance also moved more in the scanner. Specifically, no memory effects were found if mean framewise displacement was included as a covariate, analysis was limited to a subsample with mean framewise displacement below 0.1, or analysis was limited to the subsample of young adults.

### Control analyses

Various control analyses were administered to ensure that the results were not driven by nuisance factors. These included different fMRI denoising methods, additional controls for movement, and limiting analysis to a young adult subsample to prevent biases due to development and aging (see methods section for details). We also repeated the analysis exclusively on trials with correct source memory. Except for the memory effects in Study 1 (reported in the previous paragraph), the general pattern of the results remained similar in each of the control analyses (Supplementary Figures 4-9).

## Discussion

Hippocampus’ functional connectivity with the neocortex showed stable long-axis organization across resting state and memory processing, which was superseded by upregulation of connectivity with distinct regions during retrieval, but not during encoding. The aHC was more strongly connected with the temporal pole, orbitofrontal, and sensorimotor cortices, as well as parahippocampal and retrosplenial areas in the medial temporal lobes. In contrast, pHC was more strongly connected with the medial, inferior, and dorsolateral prefrontal cortices, and precunei. Similar connectivity differences along the HC long axis have been previously shown during resting statev(Adnan et al., 2016; Blessing et al., 2016; Chase et al., 2015; Libby et al., 2012; Przeździk et al., 2019; Qin et al., 2016; Robinson et al., 2016; Vos de Wael et al., 2018; Ward et al., 2014). Our results indicate that they persist during memory encoding and retrieval, confirming a previously observed phenomenon in which brain networks active during cognitive tasks also act synchronously during rest (Frank et al., 2019; Smith et al., 2009). Partial convergence with structural connectivity and gene expression patterns (Dalton et al., 2021; Vogel et al., 2020; Vos de Wael et al., 2018) further corroborates the intrinsic organization of the hippocampus. However, there are also striking differences between functional and structural connectivity, most notably the lack of direct tracts between the aHC and medial prefrontal and orbitofrontal areas (Dalton et al., 2021). HC functional connectivity is thus not fully constrained by direct structural connections and is likely mediated by other brain regions, including surrounding areas in the medial temporal lobes (Beaujoin et al., 2018; Maass et al., 2015; Ranganath and Ritchey, 2012; Ritchey et al., 2015).

In addition to the stable functional organization of the HC along its longitudinal axis, variations in connectivity strength with specific memory processes were observed for the whole HC. Preferential connectivity for encoding compared to retrieval was sparse, though there was indication of increased connectivity with sensory areas. In contrast, connectivity was stronger during retrieval than encoding with anterior temporal, medial prefrontal, inferior parietal, and parahippocampal cortices – areas previously known to be included in the supposed memory recollection network (Kim, 2010; Rugg and Vilberg, 2013). This was despite the fact that the retrieval task did have several features that were different across the tasks, for example stimulus categories, interval between encoding and retrieval, and presence of intermittent retrieval in Study 2. In other words, a core retrieval-related network emerged from the conjunction across studies, but no task-general connectome was evident for encoding. Thus, hippocampal connectivity during encoding may be more dependent on specific task requirements, in line with recent evidence showing prioritization of low-level perceptual features during encoding and higher lever conceptual features during retrieval (Linde-Domingo et al., 2019). Similarly, more widespread connectivity during retrieval than encoding speaks to increased cortical integration, potentially accompanied by decreased network modularity (Cooper and Ritchey, 2019).

We found no evidence for preferential association of aHC with encoding and pHC with retrieval. This is unexpected, as fMRI activity studies report increased aHC activity during encoding and pHC activity during retrieval (Langnes et al., 2019; Spaniol et al., 2009). Similarly, our results diverge from previously reported associations between aHC and encoding via interactions with externally oriented attentional networks (Fritch et al., 2021; Kim, 2015, 2010). Recent reports have found similar lack of specific associations with aHC and pHC with encoding and retrieval (Grady, 2020; Plachti et al., 2019). The simultaneous activation patterns in previous studies may thus reflect subsidiary cognitive processes supporting encoding and retrieval (Kim, 2020), resulting in increased fMRI activity, but not necessarily increased synchronization with the HC.

A notable exception was the relationship between the pHC and default mode network during retrieval in Study 2, on top of generally increased default mode network connectivity in both studies. While this observation agrees with some previous reports (Fritch et al., 2021; Kim, 2015), others have reported increased aHC connectivity with the default mode network compared to pHC during rest (Zheng et al., 2021, 2021). We found opposite patterns for the default mode network connectivity across the studies, further complicating the interpretation. A potential explanation for these differences is the temporal window between encoding and retrieval, which was only 90 minutes in Study 1, but 12 hours in Study 2, including sleep period, suggesting that the transformation of memory representations along the HC longitudinal axis may be affected by time-dependent consolidation (Cowan et al., 2021).

The combination of two studies with large samples is rare in task-fMRI literature, so this study stands out as a representative report of the HC-cortical connectivity. However, the combination of large sample with repeated measures design renders the interpretation of significance testing complicated. Using effect sizes instead alleviates this issue, yet there is no consensus on what constitutes a meaningful effect size in neuroscience (Marek et al., 2022), not to mention fMRI-connectivity studies (Dansereau et al., 2017). We observed generally higher effect sizes in Study 2, likely due to increased signal-to-noise ratio due to improved acquisition parameters and task design (Murphy et al., 2007). We consider the combination of statistical testing with effect sizes and the convergence across two different studies best practice to extract meaningful and reliable effects.

Turning to the limitations of the current study, the dichotomized classification of aHC and pHC using a consensus-coordinate is a simplification of HC structure and function, as converging evidence suggests a gradient along its longitudinal axis (Plachti et al., 2019; Przeździk et al., 2019). Nonetheless, a consensus cut-off point (Poppenk et al., 2013) provides a reasonable trade-off between accurate representation and analytic complexity while allowing direct comparisons with previous studies. Lastly, functional connectivity interpretation is affected by its calculation methods. When compared to implicit baseline, correlational PPI tends to yield similar results as the time-series or beta series correlation (Di et al., 2021), which may accentuate similarities between resting and task states. Further, we contrasted all task events to an implicit baseline to maximize available trials for reliable connectivity estimations, although control analysis limited to trials with correct source memory yielded similar results. As such, the observed differences between rest, encoding, and retrieval likely represent changes in cognitive states irrespective of trial-by-trial dynamics. Such state changes are expected, given that global adjustment of neural parameters is a hallmark of sustained cognition. However, the temporally sluggish fMRI-approach may be insensitive to short-term neural dynamics that eventually determine the behavioral and cognitive performance and may contribute to the lack of associations between HC-cortical connectivity and memory performance.

In conclusion, HC-cortical connectivity has distinct profiles for the aHC and pHC that are stable during resting state and memory processing. Retrieval is additionally related with upregulation of connectivity with anterior temporal, medial prefrontal, inferior parietal, and parahippocampal cortices, whereas no task-general encoding connectome emerged. These results indicate connectivity changes beyond stable network organization during memory processing.

## Materials and Methods

### Participants

Two independent samples performing different memory tasks were used in the current study. For Study 1, task data was analyzed from 514 participants (346 females and 168 males, age range 6-81 years, mean = 39, SD = 18). For Study 2, the final sample consisted of 237 participants (140 females and 97 males, age range 10-81 years, mean = 25, SD = 21). Age distribution in this sample was trimodal, with participants distributed across three age groups: children (n = 146, age range 10-16 years, mean = 12, SD = 1), young adults (n = 49, age range 20-44 years, mean = 27, SD = 5), and older adults (n = 42, age range 60-81 years, mean = 67, SD = 6). The participants in the children’s group went through a shortened study protocol (Figure 1b), excluding the resting state and final in-scanner fMRI retrieval phase. Resting state was available for all 91 adult participants.

The data was drawn from an available in-house database with screening exclusion criteria being MRI contraindications, history of disease affecting central nervous system including severe neurological or psychiatric illness and trauma, use of psychoactive drugs, and for Study 2 only, sleeping disorders. The children in Study 2 were re-invited from the subsample of Norwegian Mother Child Cohort (original subsample described in Krogsrud et al., 2014). Data was discarded due to partially missing data, deviant psychometric test results where such data was available (IQ < 85, Beck’s Depression Inventory > 20, Geriatric Depression Scale > 20, or Mini-Mental State Examination < 26), and outliers based on the average FC across all cortical regions, hippocampus seeds, and task phases +/− 3 standard deviations from the group mean.

While the wide age-ranges of these samples may offer insights into the life-span changes in HC-cortical connectivity, the initial analysis of age relationships showed complex regionally specific patterns for different cognitive states, which are beyond the scope of this report. Age and age^2^ were therefore considered as covariates of non-interest in all analyses to account for non-linear ageing effects, and control analyses on young adult subsamples were performed to prevent biases due to development and ageing.

All participants ≥12 years gave written informed consent, all participants <12 years gave oral informed consent and, and for all participants <18 years, written informed consent was obtained from their legal guardians. Ethical approval was obtained by the Regional Committees for Medical and Health Research Ethics in Norway (REK 2010/3407) and all procedures followed the Declaration of Helsinki.

### Study 1 experimental design and procedure

The procedure in Study 1 started with a 6 minutes resting state, followed immediately by an incidental memory encoding phase and then a surprise retrieval phase after approximately 90 minutes (Figure 1a,c). The details of the paradigm have been described previously (Sneve et al., 2015). Shortly, during the encoding phase, participants were presented with 100 black and white line drawings depicting everyday objects, preceded by a question either ‘Can you eat it?’ or ‘Can you lift it?’. The participants’ task was to answer the question by a button press indicating ‘yes’ or ‘no’, with the button-response mapping counterbalanced across participants. During the retrieval phase, participants were presented with all 100 items from the encoding phase and 100 new items. In each trial, between one and three questions were asked, depending on the participants’ answers. First, an item was presented on the screen, preceded by the question ‘Have you seen this item before’ (Q1). If the participant answered ‘Yes’ to Q1, a second question was asked: ‘Can you remember what you were supposed to do with the item?’ (Q2). Finally, if the participants answered ‘Yes’ to Q2, a third question was asked: ‘Were you supposed to eat it or lift it?’ (Q3), and the participants had to choose between answers ‘Eat’, and ‘Lift’. In case of ‘No’ answer to either Q1 or Q2 the trial ended without further questions. For memory analysis, a recollection index was calculated as a percentage of correct associations from all items. Approximate guessing rates for each individual was estimated by the number of incorrect answers to Q3, as the participants had indicated that they remembered the association in Q2, but then chose the wrong answer. These were subtracted from the number of correct answers before calculating the recollection index to correct for guessing (Sneve et al., 2015; Vidal-Piñeiro et al., 2021). The task presentation was controlled with E-Prime 2.0.

### Study 2 experimental design and procedure

The task in Study 2 consisted of an instructed memory encoding phase, an intermediate forced choice retrieval phase directly after encoding, and a recollection retrieval phase 12 hours after encoding (Figure 1b,d). The encoding and 12-hours retrieval were performed in the fMRI scanner. Resting state data was acquired before and after encoding, and before retrieval for 12 minutes each time. Participants under 18 years old did not return for the 12-hours retrieval sessions nor did the resting state measurement, thus only encoding fMRI data is available for this subsample. Most of the adult subsample (n = 75) completed the paradigm twice with different stimuli sets over two separate sessions, separated by a minimum of 6 days. The time of the day for encoding and the forced choice memory test was counterbalanced within-subjects, with one encoding and test session completed in the evening in one visit and the other encoding and test session completed in the morning during the other visit. The behavioral data represents the average, and the fMRI time-series data is concatenated across the two visits and resting state segments (Ness et al., 2022), except for the participants for whom only data from a single paradigm iteration was available. The trials in the encoding and final retrieval phase were separated into 2 and 3 runs, respectively, with 64 trials for each run.

During the encoding phase, participants were presented with 128 item- and face or place pairs per visit (drawn from a pool of 256 items, 8 faces, 8 places), accompanied by an auditory recording that repeated the Norwegian word for the item three times. The items consisted of real-life images of non-animate everyday items. For a single session, 4 different faces and places were used. Participants were instructed to imagine an interaction between the item and the face or place while presented on the screen, and afterwards to rate the vividness of their imagination on a scale from 1-4, in which 1 was “not vivid at all” and 4 was “very vivid”.

After the encoding phase, all participants completed a self-paced forced-choice retrieval task with 128 trials outside of the scanner. In each trial, they were presented simultaneously with the image and the recording of one of the learned items, together with all 4 faces and 4 places used in the session. Their task was to indicate by a button press, which face or place had the item been paired with during the encoding phase. All adults returned for a retrieval phase 12 hours later. Here, they were presented with 128 learned items and 64 new items (both visual and auditory) in a pseudorandomized order. The participants had to a press a button choosing between four options: (1) they had seen the item before and it was associated with a face, (2) they had seen the item before and it was associated with a place, (3) they had seen the item before, but could not remember whether it was associated with a face or place, (4) they had not seen this item before. Correct answers to (1) and (2) were considered as correct recollection in the memory analysis. Like in Study1, the recollection rates were corrected for guessing by subtracting the number of incorrect associations (wrongly choosing the answer (1) or (2)) from the number of correct associations. Task presentation was controlled with Psychtoolbox 3.

### Study 1 MRI acquisition

Imaging data were collected using a 20-channel head-neck coil on a 3T MRI (Skyra, Siemens Medical Solutions, Germany) at Rikshospitalet, Oslo University Hospital. The parameters were equivalent for resting state, encoding and retrieval phases. For each run, 43 transversally oriented slices were measured using a BOLD-sensitive T2*-weighted EPI sequence (TR = 2390 ms, TE = 30 ms, flip angle = 90°; voxel size 3×3×3 mm; FOV = 224×224 mm; interleaved acquisition; GRAPPA factor = 2). Resting state consisted of 150 volumes, each encoding run consisted of 131 volumes, while the number of volumes during retrieval was dependent on participants’ responses (207 volumes on average). Three dummy volumes were collected at the start of each fMRI run to avoid saturation effects in the analyzed data. Additionally, a standard double-echo gradient-echo field map sequence was acquired for distortion correction of the EPI images.

Anatomical T1-weighted MPRAGE images consisting of 176 sagittally oriented slices were obtained using a turbo field echo pulse sequence (TR = 2300 ms, TE = 2.98 ms, flip angle = 8°, voxel size = 1×1×1 mm, FOV = 256×256 mm).

### Study 2 MRI acquisition

Imaging data was collected using a 32-channel head coil on a 3T MRI (Prisma, Siemens Medical Solutions, Germany) at Rikshospitalet, Oslo University Hospital. The parameters were equivalent for resting state, encoding and retrieval task phases. For each run, 56 transversally oriented slices were measured using a BOLD-sensitive T2*-weighted EPI sequence (TR = 1000 ms; TE = 30 ms; flip angle = 63°; matrix = 90×90; voxel size 2.5×2.5×2.5 mm; FOV = 225×225 mm; ascending interleaved acquisition; multiband factor = 4, phase encoding direction = AP). Each encoding, retrieval, and resting state run consisted of 730 volumes. Six dummy volumes were collected at the start of each fMRI run to avoid saturation effects in the analyzed data. Sufficient T1 attenuation was confirmed following preprocessing. Additional spin-echo field map sequences with opposing phase encoding directions (anterior-posterior and posterior-anterior) were acquired for distortion correction of the EPI images.

Anatomical T1-weighted MPRAGE images consisting of 208 sagittally oriented slices were obtained using a turbo field echo pulse slices (TR = 2400 ms; TE = 2.22 ms; TI = 1000 ms; flip angle = 8°; matrix = 300×320×208; voxel size = 0.8×0.8×0.8 mm; FOV = 240×256 mm). T2-weighted SPACE images consisting of 208 sagittally oriented slices (TR = 3200 ms; TE = 5.63 ms; matrix = 320×300×208; voxel size = 0.8×0.8×0.8 mm; FOV= 256 mm×240 mm) were also obtained.

### fMRI preprocessing

fMRI was preprocessed similarly far all task phases and both studies using FMRIPREP version 1.5.3 2(Esteban et al., 2020, 2019) and Python 3.8.2. The detailed pipeline has been described previously (Ness et al., 2022). Shortly, the pipeline included BIDS-conversion, intensity nonuniformity correction, skull-stripping, susceptibility distortions correction, motion correction, co-registration with the anatomical reference using boundary-based registration with six degrees of freedom, and slice-timing correction. Distortion correction was performed using a custom implementation (https://github.com/markushs/sdcflows/tree/topup_mod) of the TOPUP technique (Andersson et al., 2003). After FMRIPREP, six motion parameters (rigid body estimation) were regressed out of the data and the individual voxel timeseries were detrended and high-pass filtered (0.008 Hz, Butterworth 5^th^ order filter). Lastly, systematic low frequency noise was removed using the *rapidtide* approach (vs. 1.9.1) (Frederick et al., 2022) with default settings, except that we did not use spatial smoothing, as signal-to-noise ratio was boosted via averaging within cortical regions instead (see next paragraph about cortical parcellation). *Rapidtide* limits the denoising to band-pass filtered data of 0.009-0.25 Hz and performs dynamic global signal regression accounting for temporal delays in systemic low-frequency oscillation propagation, avoiding artificial anticorrelations that are otherwise introduced by global signal regression (Erdoğan et al., 2016). While global signal regression is still considered controversial if several groups are compared, here we were interested in within-group regionally specific connectivity above and beyond global signal changes, necessitating its removal.

### Functional connectivity estimation

The HC was estimated based on the automatic subcortical segmentation in FreeSurfer and further divided into aHC and pHC segments as follows: for each participant, the coronal plane intersecting with the uncal apex, located at y = −21 in MNI152-space (Poppenk et al., 2013), was back-projected to the individual’s anatomy using the inverse of the nonlinear MNI-transformation established during pre-processing. HC voxels anterior/posterior to this plane were labeled anterior/posterior HC respectively. The cortex was divided into 400 regions on each participant’s native reconstructed surface based on the Local–Global Parcellation of the Human Cerebral Cortex (Schaefer et al., 2018). For the resting state, functional connectivity of the two HC seeds (aHC and pHC) with each cortical region was estimated by correlating their BOLD timeseries. The task functional connectivity was calculated by correlational psychophysiological interaction analysis (cPPI; Fornito et al., 2012), resulting in symmetrical undirected matrices for aHC and pHC connectivity with each of the 400 cortical regions using all task events as described previously (Capogna et al., 2022). That is, the psychological timeseries were constructed as boxcar functions, reflecting 2 s or 5 s events from the start of the trial for Study 1 and 2 respectively, matching the stimulus duration. The PPI terms were constructed by first deconvolving BOLD timeseries from each region (Gitelman et al., 2003), then multiplying these point-by-point with the psychological event timeseries, and finally re-transforming these to BOLD through convolution with a canonical 2-gamma HRF. Finally, the cPPI matrices were constructed by partial Pearson’s correlation analysis between the constructed PPI terms of the HC and each cortical region, controlling for either regions’ BOLD timeseries and the HRF-convolved psychological timeseries. The resulting correlation coefficients were Fisher-transformed to z-values and averaged over left and right HC seed regions. For the default mode, dorsal-attentional, salience and frontoparietal control networks, the z-values were averaged across the regions belonging to each network, as defined by Schaefer et al., (2018) 7-network parcellation (Yeo et al., 2011). The cPPI calculation was done using Matlab 2017a, spm 12.0, and gPPI toolbox 13.1.

### Statistical analysis

Whole brain analyses were performed using mixed effects linear models (function *fitlme*) in Matlab (version 2018b) separately for resting state and task data. First, functional connectivity at each HC-cortical pair was predicted by HC segment (aHC vs pHC), controlling for age, age^2^, and sex, and including participant ID as a random variable. For task data, this model was expanded by adding task phase (encoding vs retrieval) and its interactions with the HC segment. Lastly, memory performance and corresponding interactions were added as additional predictors. The resulting contrast effects were corrected for multiple comparisons using false discovery rate correction (Benjamini and Hochberg, 1995) with a threshold of p<0.05. The conjunction across cognitive states and/or studies was considered significant if the fdr-corrected effects were significant in all states/studies included in the analyses (Nichols et al., 2005). The contrast t-values with corresponding degrees of freedom were transformed into Cohen’s d using the *effectsize* package(Ben-Shachar (@mattansb) et al., 2022) and visualized using the *ggseg* package (Mowinckel and Vidal-Piñeiro, 2020) in R version 3.6.2. The correlation coefficients’ significance of either the raw values or t-contrast maps across different studies and cognitive states was tested via permutation analyses controlling for spatial autocorrelation using the BrainShmash python package (Burt et al., 2020).

Network analysis and visualization were performed using R version 3.6.0 and package collections *tidyverse* (Wickham et al., 2019), and *lme4* (Bates et al., 2022). Mixed effects linear models with planned contrasts were set up with average HC-network connectivity as an outcome variable, and HC segment (aHC vs pHC), cognitive state (rest vs task contrast: rest = 1, encoding = −0.5, retrieval = −0.5; encoding vs retrieval contrast: rest = 0, encoding = 1, retrieval = −1), controlling for age and sex, and participant ID as a random variable.

### Control analyses

As fMRI connectivity is notoriously susceptible to denoising methods (Ciric et al., 2017), we repeated the whole brain linear models with two different denoising methods. First, we used six motion estimates and two physiological time-series (one-step eroded white matter and cerebrospinal fluid masks), combined with static global signal regression as nuisance parameters to reduce the effect of motion. Second, we repeated the analyses using the independent component analysis based method ICA-AROMA (Pruim et al., 2015), again with mean white matter and cerebrospinal fluid timeseries as nuisance regressors. Of note, several regions with negative connectivity were observed for the denoising method using static global signal regression in Study 1, but given that these were lacking in Study 2 and when different denoising methods were used, these were likely spurious (Erdoğan et al., 2016). Similarly, the aHC and pHC effects results diverged initially using ICA-AROMA from both other methods, yet this was caused by global signal variations during different cognitive states. When average connectivity across all regions was included as a covariate in the linear models, the patterns of connectivity were again similar to the two other methods.

In-scanner movement tends to correlate with functional connectivity estimates despite rigorous data cleaning and de-noising (Ciric et al., 2017), which we also found in our data (correlation coefficients between average framewise displacement (fd) and mean connectivity ranging between 0.13 and 0.47 across studies and cognitive states). We repeated the whole brain linear models using mean fd as a covariate, and once more using only a subsample of participants with fd < 0.1 (rest n =265, task n = 228). Note that the latter analysis was limited to Study 1, since >90% of the participants in Study 2 had fd < 0.1 to start with.

A possible limitation for secluding core memory effects is that our analysis was run on all task data. Therefore, we repeated the analysis on datasets in which we included only trials with correct source memory performance. Despite lower signal-to-noise ratio due to decreased number of trials in this analysis, the results did not change.

Lastly, while we consider the large representative age range of our studies a strength, the inclusion of children and elderly may bias the results towards developmental or aging effects, respectively. We therefore repeated the linear models on subsamples of young adults from age 18-40 (Study 1 rest n = 268, task n = 266; Study 2 rest n = 48, task n = 48).

## Acknowledgments

The studies were supported by European Research Council (725025 and 286634) and Norwegian Research Council (262453).

## Data availability

All group-level connectivity maps are available at the dedicated project at Open Science Framework (https://osf.io/834hz/). The raw MRI data may be available upon reasonable request, given appropriate ethical, data protection, and data-sharing agreements. Requests for the raw MRI data can be submitted to the principal investigator (Prof. Anders Fjell, University of Oslo). Individual-level data availability may be restricted as participants have not consented to publicly share their data.

## Code availability

Statistical analysis scripts (together with simulated data) are available at the dedicated project at Open Science Framework (https://osf.io/834hz/).

