## Supplementary Materials for "Hippocampal-cortical functional connectivity during memory encoding and retrieval"

### during memory encoding and retrieval

#### Supplementary materials

Liisa Raud

2022-08-11

#### Raw HC-cortical connectivity

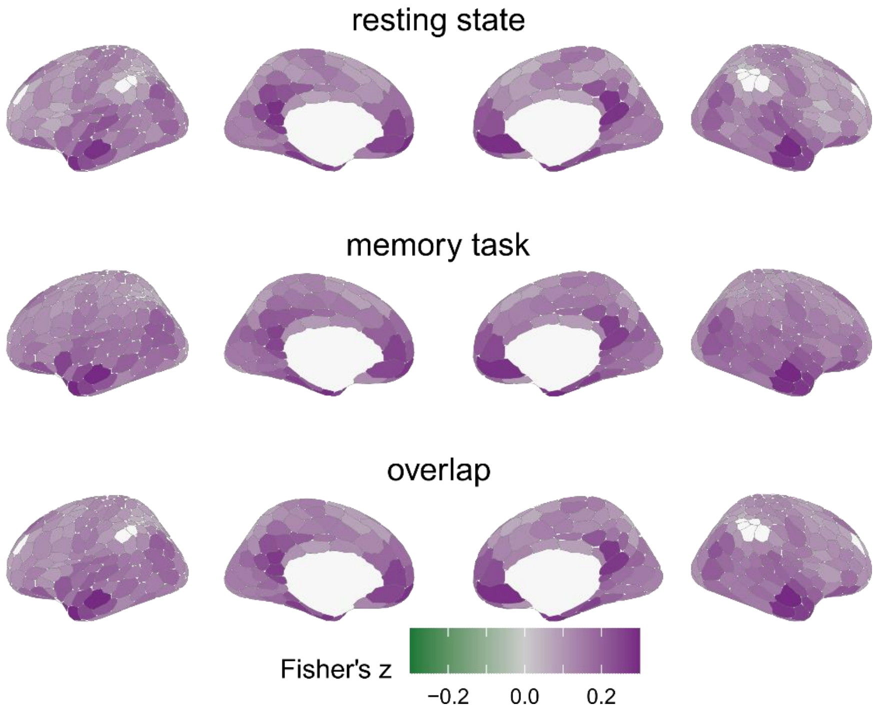

**Figure 1. HC-cortical functional connectivity for resting state, task state, and their overlap**. The values are averaged across the aHC and pHC, the two studies, and encoding and retrieval states for the task connectivity. Overlap plot indicates averaged z-values across both states (rest and task) and studies. Values are masked for significant conjunctions where intercepts in both studies and states (linear model predicting connectivity from HC segment, controlling for sex, de-meaned age and $age^2$) were significant at $p < 0.05$ (fdr-corrected).

#### Memory effects

Linear mode for each HC-cortical ROI pair: connectivity = 1 + hippocampus segment (anterior/posterior) * task phase (encoding/retrieval) * memory + age + age2 + sex + 1|ID

##### Resting state

There were no memory main effects in resting state in either study. There were limited interactions between hippocampus segment and memory (3 parcels in Study 1: left retrosplenial cortex, left somatomotor cortex, left temporal pole; 4 parcels in Study 2: bilateral insulae, right posterior medial prefrontal cortex), which did not overlap across the studies.

##### Task state

There were no memory main effects in either study, but due to extensive interactions with hippocampus ment (Figure 2), the analyses are repeated for anterior and posterior hippocampus separately. Most regions with previous interaction effects were not significant in those models, but there were negative associations with memory (i.e. higher connectivity related to worse memory performance) with lateral and medial temporal cortices, medial prefrontal, and medial parietal cortices, limited to Study 1 (Figure 3). There were no significant memory effects in the follow-up analysis for Study 2.

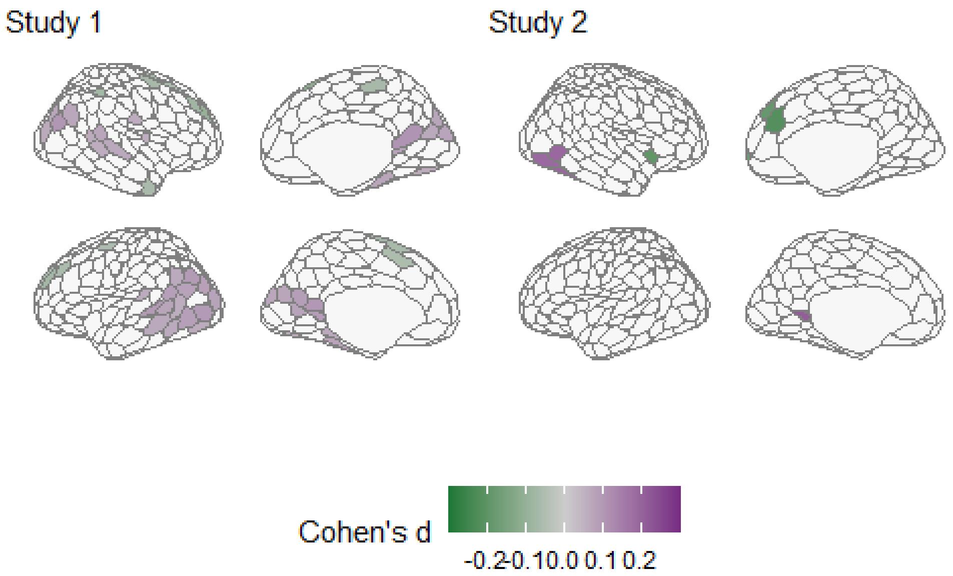

**Figure 2. Memory and hippocampus subfield interactions.**

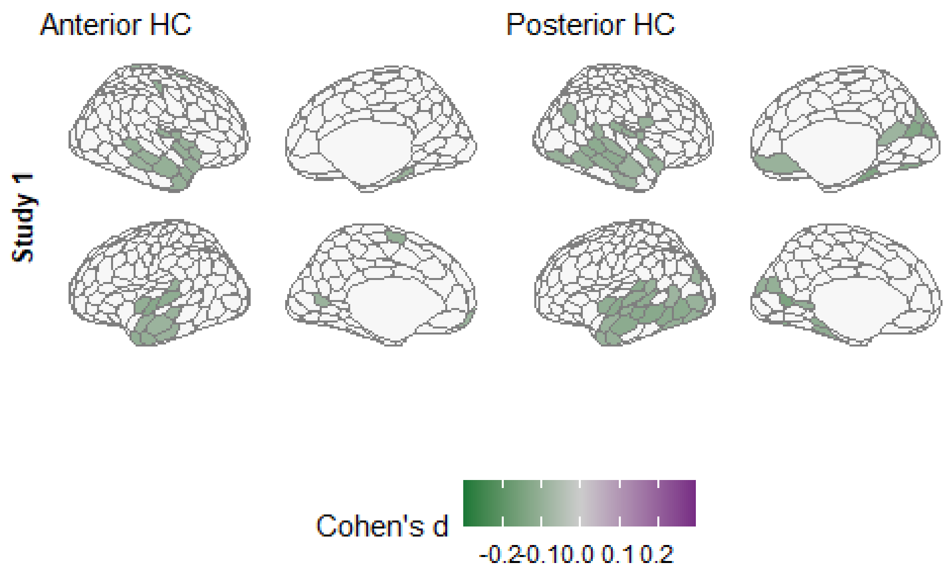

**Figure 3. Memory main effects in Study 1.**

#### Control analyses for memory effects

The interaction effects of memory and hippocampus segment (Figure 2) disappeared in the following control analyses: 1. analyses limited to a subsample with average framewise displacement below 0.1 2. analyses limited to a subsample of young adults

While the interaction effects initially were similar to the original results (Figure 2) for the control analysis with average framewise displacement as a covariate, the main effects in the follow-up analyses (Figure 3) were no longer significant.

#### Control analyses

##### Movement: average framewise displacement as a covariate

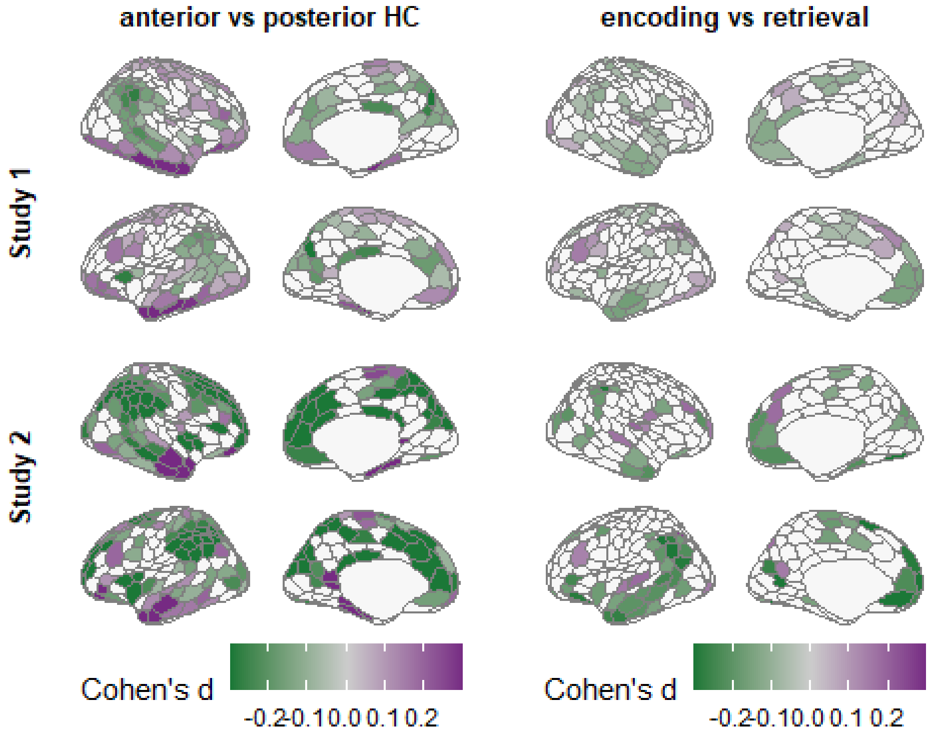

**Figure 4. Main effects with movement covariate.**

#### Movement: subsample of participants with average framewise displacement below 0.1

Note that this analysis is limited to Study 1, since >90% of participants in Study 2 had low movement parameters to begin with.

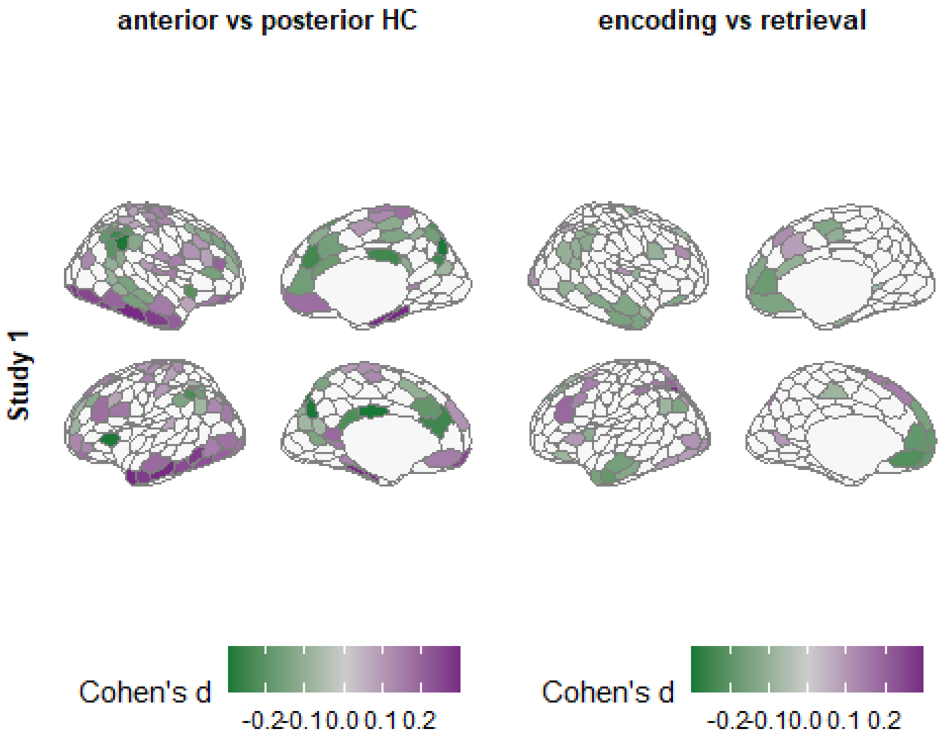

**Figure 5. Main effects on low-movement subsample.**

#### Developmental and ageing effects: limiting the analysis to a subsample of young adults

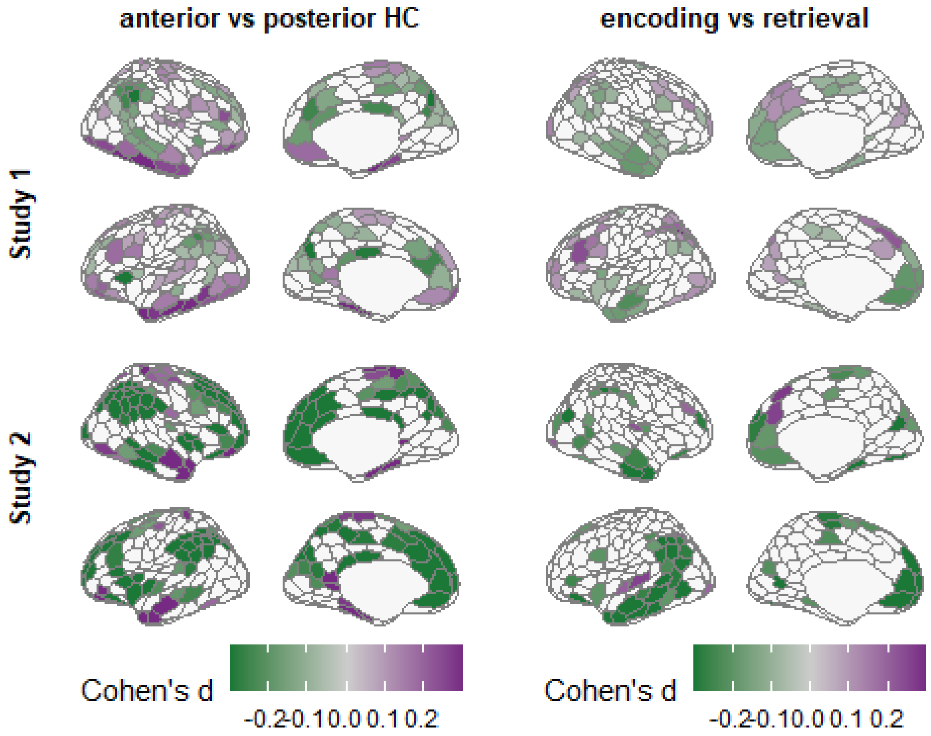

**Figure 6. Main effects on young adults subsample.**

#### Denoised

fMRI denoising done using six motion estimates and two physiological time series (one-step eroded WM and CSF masks), combined with static global signal regression as nuisance parameters.

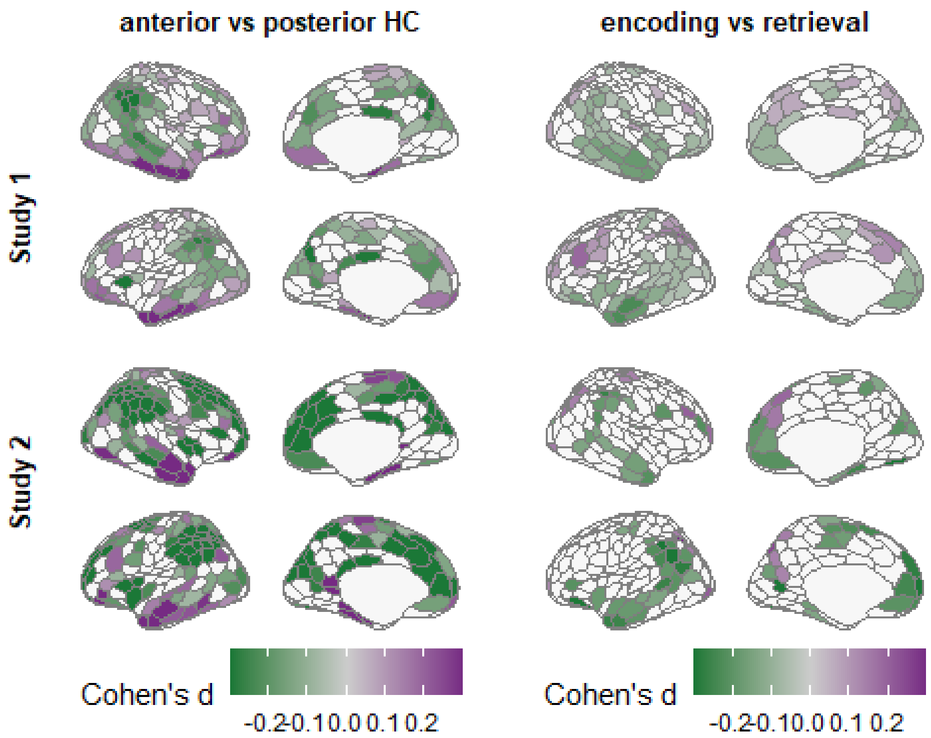

Figure 7. Main effects with different denoising.

#### ICA-aroma

fMRI denoising done using ICA-AROMA and two physiological time series (one-step eroded WM and CSF masks) and average connectivity across all ROIs as nuisance parameters.

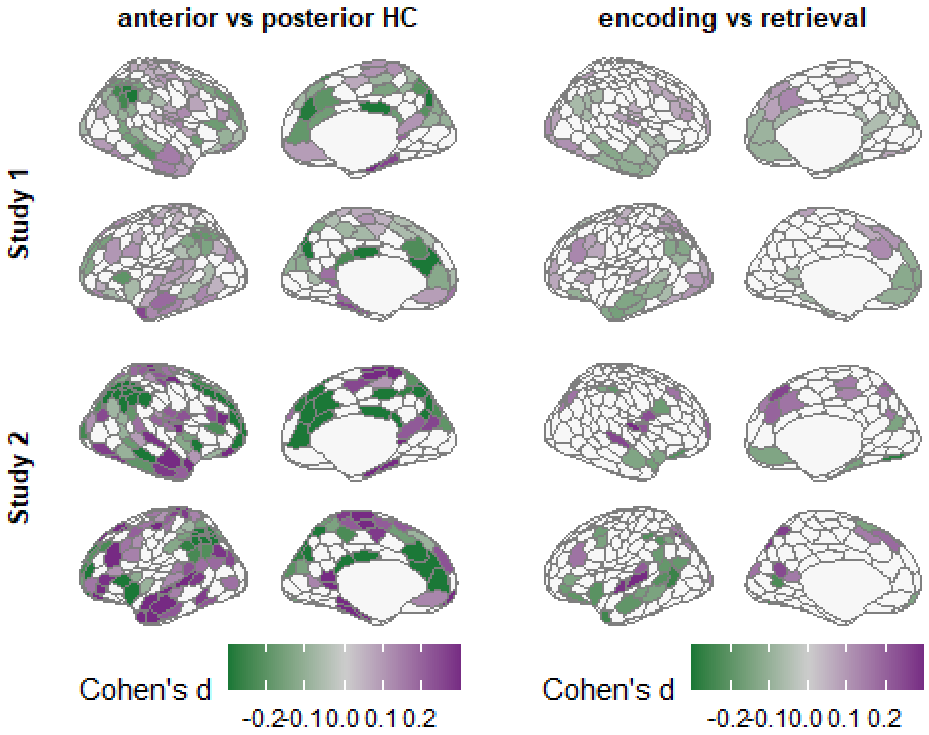

Figure 8. Main effects with ICA-AROMA.

#### Limiting analysis on trials with correct source memory performance

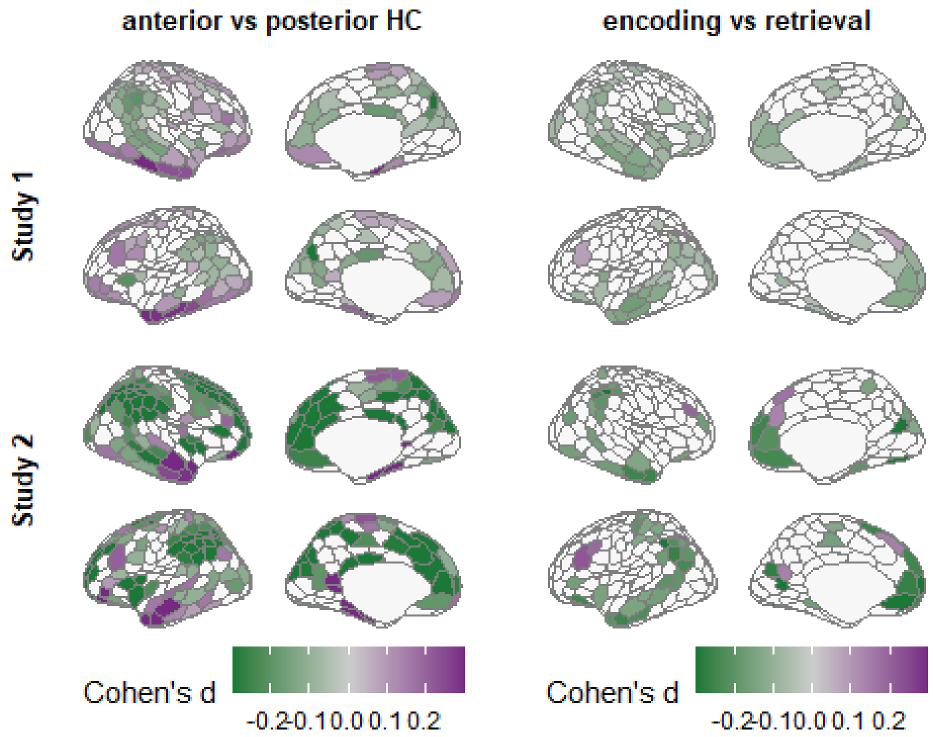

**Figure 9. Main effects on correct source memory trials.**
